# Antiviral Type III CRISPR signalling via conjugation of ATP and AdoMet

**DOI:** 10.1101/2023.06.26.546636

**Authors:** Haotian Chi, Ville Hoikkala, Sabine Grüschow, Shirley Graham, Sally Shirran, Malcolm F White

## Abstract

CRISPR systems are widespread in the prokaryotic world, providing adaptive immunity against mobile genetic elements (MGE) ^1, 2^. Type III CRISPR systems, with the signature gene *cas10*, use CRISPR RNA (crRNA) to detect non-self RNA, activating the enzymatic Cas10 subunit to defend the cell against MGE either directly, via the integral HD nuclease domain ^3–5^ or indirectly, via synthesis of cyclic oligonucleotide (cOA) second messengers to activate diverse ancillary effectors ^6–9^. A subset of type III CRISPR systems encode an uncharacterised CorA-family membrane protein and an associated NrN family phosphodiesterase predicted to function in antiviral defence. Here, we demonstrate that the CorA associated type III-B (Cmr) CRISPR system from *Bacteroides fragilis* provides immunity against MGE when expressed in*E. coli*. However, *B. fragilis* Cmr does not synthesise cOA species on activation, instead generating a previously undescribed signalling molecule, SAM-AMP (3’-adenylyl-AdoMet) by conjugating ATP to S-adenosyl methionine via a phosphodiester bond. Once synthesised, SAM-AMP binds to the CorA effector, presumably leading to cell death by disruption of the membrane integrity. SAM-AMP is degraded by CRISPR associated phosphodiesterases or a SAM-AMP lyase, providing an “off switch” analogous to cOA specific ring nucleases ^10^. SAM-AMP thus represents a new class of second messenger for antiviral signalling, which may function in different roles in diverse cellular contexts.

---

*Bacteroides spp.* are Gram-negative, anaerobic bacteria that constitute a significant portion of the human gut microbiome ^11^. The *Bacteroidales* are host to the most widespread and abundant phage found in the human digestive system, CrAssphage ^12^. *B. fragilis* is an opportunistic pathogen, responsible for >70 % of *Bacteroides* infections ^13^. Bioinformatic analyses has revealed the presence of three CRISPR types: I-B, II-C and III-B, in *B. fragilis* strains, with the type III-B system most common ^14^. Sequence analysis shows that *B. fragilis* Cas10, the main enzymatic subunit of type III effectors, lacks an HD nuclease domain but has an intact cyclase domain, similar to the *Vibrio metoecus* Cas10 ^15^. This suggests that the system functions via cyclic oligoadenylate (cOA) signalling to associated ancillary effectors. In *B. fragilis* and more generally in the CFB bacterial phylum these type III CRISPR systems are strongly associated with an uncharacterised gene encoding a divergent member of the CorA-family of divalent cation channel proteins ^16, 17^ (Fig. 1a). The CRISPR associated CorA proteins have not been studied biochemically but are predicted to consist of a C-terminal membrane spanning helical domain fused to a larger N-terminal domain with a unique fold. To investigate this further, we first generated a phylogenetic tree of all Cas10 proteins and identified those associated with a gene encoding the CorA protein. Three phylogenetically distinct clusters of CorA-associated type III CRISPR systems were apparent, with the largest (CorA-1) associated with type III-B systems (Fig. 1a).

**Fig. 1.**
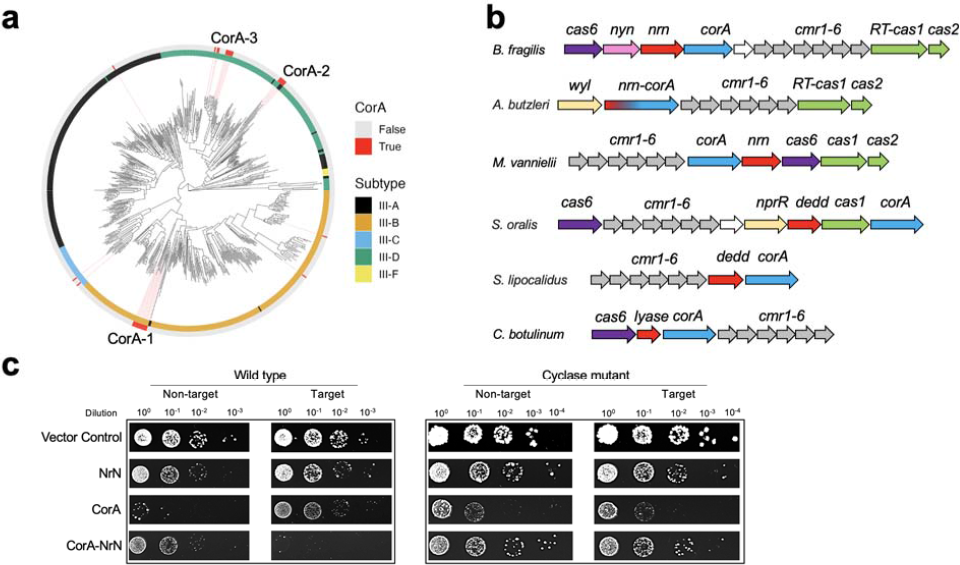
Type III CRISPR systems with a CorA effector. **a,** Phylogenetic tree of Cas10 proteins from type III CRISPR-Cas systems of complete bacterial and archaeal genomes, colour coded by subtype ^18^. Red bars on the outer ring indicate systems associated with a CorA family effector protein. Three main clusters of CorA-associated Cas10s are observed, labelled CorA-1, −2 and −3. **b,** Genome context and effectors of selected type III-B CRISPR systems with a *corA* gene (cluster CorA-1). The type III-B *cas* genes *cmr1-6* are shown in grey, with *cas6* in purple and the adaptation genes *cas1* (or a gene encoding a fused reverse transcriptase-cas1 protein) and *cas2* in green. The putative membrane channel protein is encoded by the *corA* gene (blue), which is adjacent to or fused with the genes encoding PDEs NrN or DEDD (red). In *Clostridium botulinum*, the PDE is replaced with a predicted SAM lyase. The *wyl* and *nprR* genes encode predicted transcriptional regulators. **c,** Plasmid Challenge Assay. *E. coli* BL21 Star cells expressing *B. fragilis* Cmr (wild-type or cyclase defective variant) programmed with target (tetR) or non-target (pUC19) crRNA species were transformed with a pRAT plasmid that expressed the NrN and/or CorA proteins and carried a tetracycline resistance gene. Resistance was only observed when a targeting crRNA, active cyclase and both effector proteins were present. Raw data available in Supplementary figure 1.

The genomic context of CorA-containing type III CRISPR loci from cluster CorA-1 (Fig. 1b) reveals the *corA* gene is typically found next to a gene encoding a phosphodiesterase (PDE) – the DHH-family nuclease NrN in the case of *B. fragilis* and *Methanococcus vanielii,* and a DEDD-family nuclease in the case of *Streptococcus oralis* and *Syntrophothermus lipocalidus*. In the genome of *Aliarcobacter butzleri* and related species, the *nrn* and *corA* genes are fused, suggestive of a close functional relationship. The closest predicted structural matches for *B. fragilis* NrN are to the pGpG-specific PDE PggH from *Vibrio cholerae*, which plays a role in the turnover of the cyclic nucleotide c-di-GMP ^19^ and the GdpP PDE from *Staphylococcus aureus*, which degrades pApA molecules as a component of c-di-AMP signalling systems ^20^. Analysis of the DEDD protein suggests structural matches to RNaseT ^21^, Oligoribonuclease (ORN) ^22^ and the mammalian REXO2 protein which degrades linear RNA and DNA dinucleotides ^23^. Thus, the CRISPR-associated NrN and DEDD proteins appear homologous to protein families that degrade small linear RNA and DNA species. Intriguingly, in some CorA-containing type III systems including *Clostridium botulinum*, the PDE is replaced by a protein predicted to resemble a family of phage SAM lyase enzymes involved in evasion of host immune systems (Fig. 1b) ^24, 25^.

## The *B. fragilis* Cmr system provides immunity against MGE in *E. coli*

To investigate the activity of the *B. fragilis* type III CRISPR system, two plasmids were constructed. Plasmid pBfrCmr1-6, built using Gibson assembly ^26^, expresses synthetic versions of the codon optimised *cmr*1-6 genes and plasmid pBfrCRISPR encodes Cas6 and a mini-CRISPR array (Extended Data, Fig. 1a). We expressed the complex in *E. coli* with a targeting (pCRISPR-Tet) or non-targeting (pCRISPR-pUC) crRNA and challenged cells by transformation with a pRAT-Duet plasmid expressing one or both of the CorA and NrN effector proteins (Fig. 1c). The pRAT-Duet plasmid also had the *tetR* gene for activation of *B. fragilis* Cmr carrying the targeting crRNA. Cells were transformed with the pRAT-Duet vectors and grown in the presence of tetracycline to select for transformants. We included vectors expressing wild-type and cyclase defective (Cas10 D328A:D329A variant) Cmr for comparison. We previously used this experimental design to investigate the *V. metoecus* Cmr system ^15^. In conditions where the Cmr system was activated and had the required ancillary effector proteins, lower numbers of colony forming units (cfu’s) were expected. The vector control (no effectors) served as a baseline for transformation. When only the NrN effector was present, no reduction in cfu’s was observed, suggesting no active targeting. When only the CorA protein was expressed, reduced cfu’s were observed in both target and non-target conditions, for both the wild-type and Δcyclase Cmr, suggesting some toxicity of the CorA protein. When both the CorA and NrN effectors were expressed, immunity (strongly reduced cfu) was observed only for the wild-type Cmr system with *tetR* targeting. Immunity was lost when a variant of NrN with mutation of the DHH active site motif (D85A:H86A:H87A), or when the CorA protein was truncated to remove the trans-membrane domain (Extended Data, Fig. 1b).

These data suggest that the *B. fragilis* Cmr system is functional in *E. coli* and requires the activity of the Cas10 cyclase domain and the presence of both effector proteins. The toxicity of CorA appears to be reduced by the presence of the NrN protein, regardless of activation of the type III system, suggestive of a strong functional link. Intriguingly, the type III-B complex from *Mycobacterium tuberculosis,* which synthesises cA_3-6_ *in vitro* ^27^, did not provide immunity when combined with the NrN and CorA effectors, hinting at a non-canonical activation mechanism (Extended Data, Fig. 1c).

## CRISPR RNA processing and target RNA degradation

Co-transformation of the expression plasmids into *E. coli* strain BL21 (DE3) allowed the expression of the *B. fragilis* Cmr effector and purification by immobilised metal affinity and size exclusion chromatography (Extended Data, Fig. 2a). We also purified *B. fragilis* Cas6 individually using the same chromatography steps. We first confirmed that Cas6 processed crRNA in the expected manner. The recombinant Cas6 enzyme cleaved synthetic FAM-labelled crRNA at the base of a predicted hairpin with a 2 bp stem, reminiscent of *Methanococcus maripaludis* Cas6b ^28^. This generates a canonical 8 nt 5’-handle, (Extended Data, Fig. 2b). Cleavage of an *in vitro* transcript composed of 2 repeats flanking one spacer generated the expected set of reaction products, culminating in a processed crRNA of 72 nt (Extended Data, Fig. 2c). To investigate the composition of the crRNA present in the effector complex purified from *E. coli*, we isolated and labelled the crRNA using γ-^32^P-ATP and polynucleotide kinase. This revealed 3 major crRNA species differing in length by 6 nt (Extended Data, Fig. 3a, b). These products correspond to 3’ end trimming of the crRNA to remove the repeat derived sequence and likely reflect effector complexes that differ in the number of Cas7 subunits and thus length of backbone, as has been seen for other type III systems (reviewed in ^29^).

**Fig. 2.**
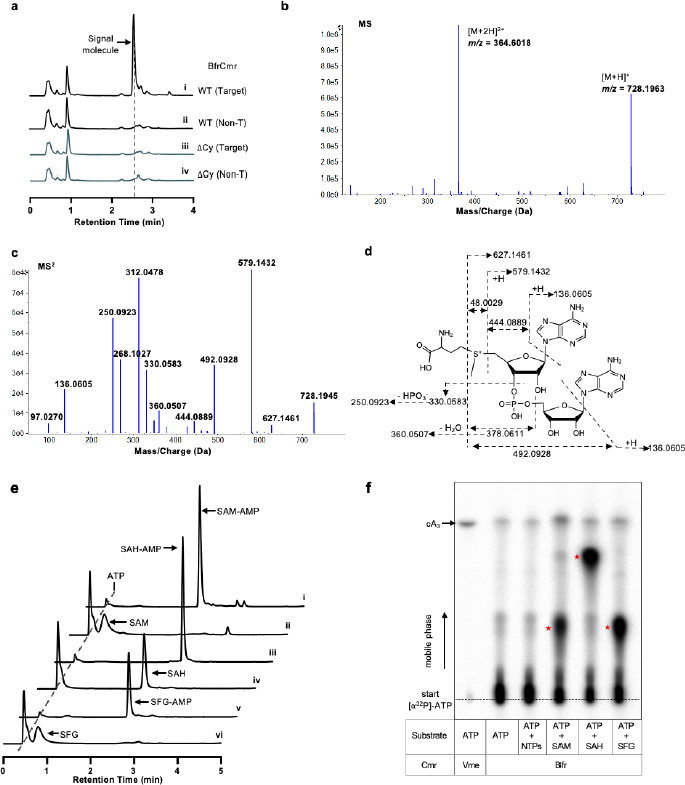
Identification of SAM-AMP from cells harbouring the activated Cmr complex. **a,** HPLC analysis of *E.coli* extracts expressing the wild type or mutant (ΔCy) *B. fragilis* Cmr system with target or non-target crRNA. The putative signal molecule was only observed for the activated system (trace i). **b,** Characterisation of the extracted signal molecule by LC-MS in positive mode. [M+H]^+^ and [M+2H]^2+^ are two different ionization forms. **c,** MS/MS analysis of the signal molecule with m/z 728.1963. **d,** The proposed structure of the signal molecule, whose fragmentation pattern is shown by dotted arrows. **e,** HPLC analysis of compounds synthesised by the purified wild type *B. fragilis* Cmr complex *in vitro*. Cmr synthesises the signal molecule 3’-adenylyl-AdoMet (SAM-AMP) from ATP and S-adenosyl methionine (trace i). Cmr also accepts S-Adenosyl-L-homocysteine (SAH) and sinefungin (SFG) as substrates (traces iii and v, respectively). Traces ii, iv and vi are control reactions. **f,** TLC analysis of *in vitro* reaction products. SAM, SAH and sinefungin plus ATP yielded radioactive products (red stars) but ATP alone did not. cA_3_ generated by wild type *V. metoecus* Cmr complex^15^ is shown for comparison. Uncropped HPLC and TLC data are available in Supplementary Fig. 2.

**Fig. 3.**
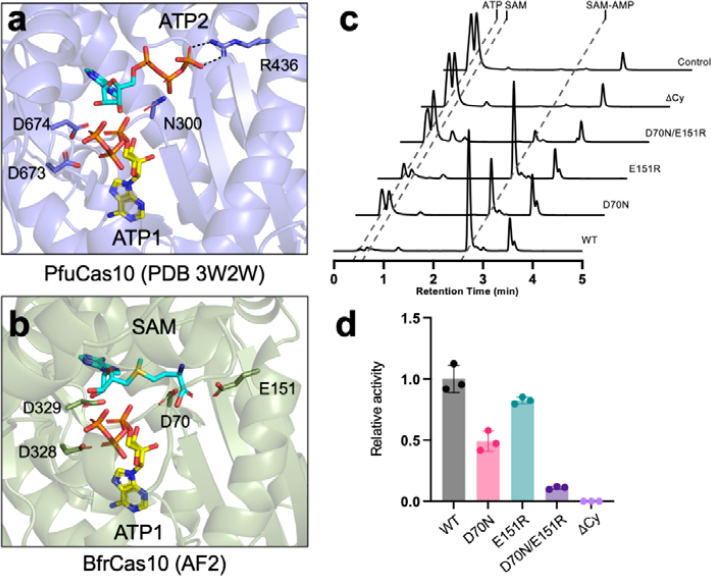
Changes in the active site of Cas10 synthesising SAM-AMP. **a,** The crystal structure of the *P. furiosus* (Pfu) Cas10 subunit with 2 ATP molecules bound ^34^. Side chains for the two metal binding aspartate residues of the “DD” motif, together with residues N300 and R436 that interact with ATP2, are shown. **b,** Equivalent view of the AF2 model of the *B. fragilis* Cas10 structure with ATP1 from the PfuCas10 structure and ATP2 replaced by SAM. The precise conformation and position of SAM is unknown. The conserved acidic residues D70, E151, D328 and D329 are shown. **c,** *In vitro* SAM-AMP synthase activity of wt and variants *B. fragilis* Cmr, analysed by HPLC following incubation of 2 µM Cmr with 0.5 mM ATP and SAM for 30 min. Raw HPLC data are available in Supplementary Fig. 3. **d,** Relative SAM-AMP synthase activity of Cmr variants. Three independent experiments were carried out, with the mean and standard deviation shown.

We proceeded to test for cleavage of target RNA bound to the crRNA in the effector. 5’-end labelled target RNA was cleaved at four positions with 6 nt spacing, corresponding to the placement of the Cas7 active sites in the backbone ^30^. Cleavage was extremely rapid and was essentially complete after 2 min, the first time point (Extended Data, Fig. 3c. As these sites interconvert and site 1 is farthest from the 5’ label, it was only observed for the Cmr4/Cas7 D27A variant, which cleaves target RNA more slowly (Extended Data, Fig. 3d). We also observed cleavage of target RNA at the boundary of the crRNA:target RNA duplex. This activity, which has not been observed for other type III systems, appeared to be due to the Cmr4 subunit, as it was not observed for the D27A variant. As target RNA cleavage has been shown to correlate with the deactivation of the Cas10 subunit ^8, 31^, this suggests that the Cmr complex remains active for a very short time after target RNA binding. This groups *B. fragilis* Cmr together with the type III effectors from *Streptococcus thermophilus* and *Thermotoga maritima*, which cleave target RNA rapidly ^5, 31^. In contrast, the type III systems from *S. solfataricus* and *V. metoecus* have much slower RNA cleavage kinetics ^8, 15^.

## *B. fragilis* Cmr synthesises a novel second messenger, SAM-AMP

As *B. fragilis* Cmr lacks an HD nuclease domain in the Cas10 subunit, immune function is expected to be mediated by the cyclase domain active site via generation of nucleotide second messengers. However, although the system provided cyclase-dependent immunity *in vivo*, activation of the wild-type Cmr *in vitro* resulted in very low yields of any observable product when incubated with ATP, in contrast to the Cmr complex from *Vibrio meteocus*, which synthesises cA_3_ ^15^ (Fig. 2f). These data hinted at the possibility that a vital component was missing in the *in vitro* assays. Accordingly, we activated *B. fragilis* Cmr in *E. coli* using a plasmid to express target RNA and then processed cell lysates to allow isolation of nucleotide products. These were purified and analysed by HPLC. A significant peak was observed following HPLC of extracts with activated Cmr, which was absent in the absence of target RNA or when the cyclase activity was knocked out by mutagenesis (ΔCy) (Fig. 2a). Mass spectrometry yielded a m/z value of 728.19 for the positive ion, a value which did not match any known cyclic nucleotide or indeed any other previously characterised metabolite (Fig. 2b). To identify the product, we fragmented the purified molecule using tandem MS/MS. This allowed the identification of fragments characteristic of AMP and methionine (Fig. 2c). Further examination suggested that the molecule under study was *S*-adenosyl methionine (AdoMet, SAM) adenylated on the ribose moiety (Fig. 2d), a molecule we henceforth term SAM-AMP. There is no published description of SAM-AMP, either from chemical or enzymatic synthesis routes, suggestive of a new, previously undiscovered class of signalling molecule.

To confirm that *B. fragilis* Cmr synthesised SAM-AMP, we reconstituted the reaction *in vitro* with ATP and AdoMet, analysing reaction products by HPLC and TLC (Fig. 2e, f). We observed SAM-AMP production when AdoMet and ATP were present *in vitro*. Substitution of AdoMet by S-adenosyl homocysteine (SAH) or the AdoMet analogue sinefungin ^32^, which differ on the sulfur centre, were also conjugated with ATP by Cmr (Fig. 2e, f; Extended Data, Fig. 4c). No significant products were observed in the presence of ATP or all four ribonucleotides. The synthesis of SAM-AMP and SAH-AMP by *B. fragilis* Cmr were consistent with rapid, multiple-turnover kinetics that were essentially complete with the first 2 min of the reaction (Extended Data, Fig. 4a, b). The observation of only SAM-AMP, and not SAH-AMP, in *E. coli* cell extracts is likely due to the fact that the concentration of SAM is much higher than SAH in *E. coli* (0.4 mM versus 1.3 µM) ^33^. Overall, these data provide strong evidence that the *B. fragilis* Cmr system generates a previously undescribed conjugate of AdoMet and ATP, rather than cOA.

**Fig. 4.**
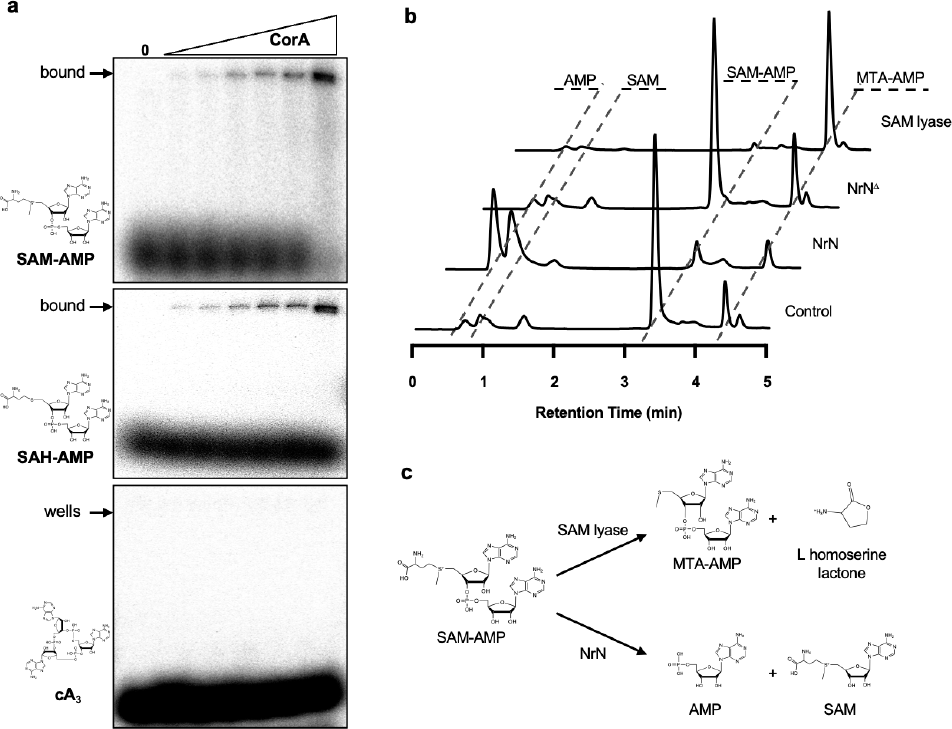
Binding and degradation of SAM-AMP by ancillary CRISPR proteins. **a,** CorA binds SAM-AMP and SAH-AMP, but not cA_3_ (1 µM ^32^P-labelled ligand incubated with 0, 0.0625, 0.125, 0.35, 0.75, 1.5, 3.3 µM CorA), illustrated by acrylamide gel electrophoresis and phosphorimaging. Uncropped gels are available in Supplementary Fig. 3b. **b,** NrN specifically degrades SAM-AMP to SAM and AMP. HPLC analysis of samples in which purified SAM-AMP was incubated with NrN and NrN^Δ^, an inactive variant (D85A/H86A/H87A). *C. botulinum* lyase degrades SAM-AMP to generate methylthioadenosine (MTA) and L-homoserine lactone (not UV visible). Small amounts of MTA are present in the SAM-AMP sample purified from *E. coli*. Uncropped HPLC traces are available in Supplementary Fig. 3c. **c,** Schematic representation of the reactions catalysed by NrN and SAM-AMP lyase.

Since Cas10 family enzymes synthesise 3’-5’ phosphodiester bonds ^6, 7^, we considered it highly likely that AdoMet was fused to AMP at the 3’ position on the ribose ring, but the MS data could not rule out a 2’-5’ phosphodiester bond. To resolve this, we incubated SAM-AMP and SAH-AMP with nuclease P1, which is specific for 3’-5’ phosphodiester bonds. Both molecules were broken down into their constituent AMP and SAH/SAM components, consistent with a 3’-5’ phosphodiester linkage (Extended Data, Fig. 5).

**Fig. 5.**
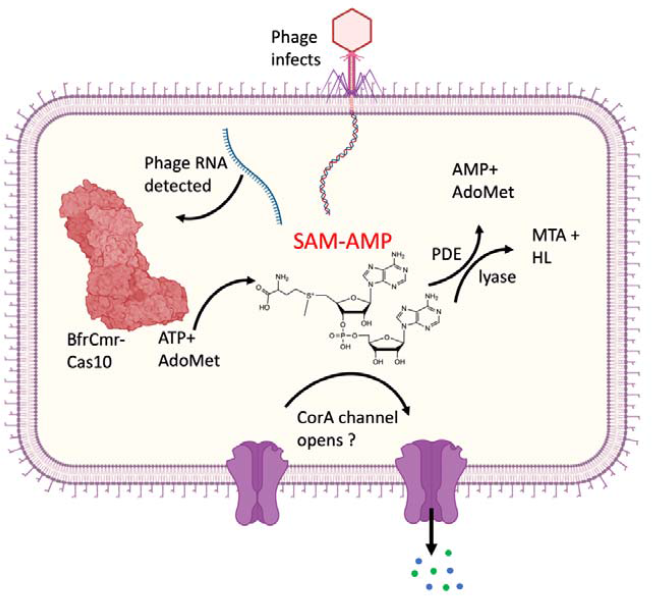
Model of the SAM-AMP immune signalling pathway. Transcription of the infecting phage genome activates the *B. fragilis* Cmr complex, resulting in synthesis of the SAM-AMP second messenger. SAM-AMP binds to the CorA membrane protein, resulting in the opening of a pore that disrupts the host membrane to combat infection. SAM-AMP is degraded by specialised PDE enzymes that hydrolyse the phosphodiester bond, generating AMP and AdoMet or lyases that target the methionine moiety, generating MTA and homoserine lactone. These enzymes likely deactivate the signalling molecule to reset the system once phage have been eliminated. Created with BioRender.com.

The crystal structure of *Pyrococcus furiosus* Cas10-Cas5 bound to two ATP molecules ^34^ shows one ATP in the “donor” ATP1 site next to the GGDD cyclase catalytic motif and another in the “acceptor” site (Fig. 3a). For the enzymes that synthesise SAM-AMP, S-adenosyl methionine must bind in the “acceptor” ATP2 binding site, next to ATP1 in the donor site ^35^. This arrangement would allow nucleophilic attack from the 3’-hydroxyl group of SAM to the α-phosphate of ATP1, resulting in the formation of a 3’-5’ phosphodiester bond linking SAM with AMP, and the release of pyrophosphate. The reaction chemistry is essentially the same as that which takes place in canonical type III CRISPR systems which synthesise cyclic oligoadenylates ^35^. The major difference is that the triphosphate of ATP2 in the acceptor site is replaced by the methionine moiety of SAM, resulting in a change in local charge from −3 to +1 in the ligand, raising the possibility that Cas10s binding AdoMet will have a less basic binding site in this area. Examination of sequence conservation in the Cas10s associated with a CorA-1 cluster (Extended Data, Fig. 6) revealed the presence of two absolutely conserved acidic residues, D70 and E151. Modelling of the *B. fragilis* Cas10 structure using Alphafold2 ^36^ places these two residues in the vicinity of the methionine moiety of AdoMet in the acceptor site (Fig. 3b). D70 occupies the position equivalent to N300 in PfuCas10, which neighbours the β-phosphate of the acceptor ATP2 ligand, while E151 is in a similar position to R436, which forms a bidentate H-bond with the γ-phosphate (Fig. 3a). We created variants of Cas10 with mutations D70N, E151R and the double mutant, which were expressed and purified as for the wild-type protein (Extended Data, Fig. 2a), and assessed for their ability to synthesise nucleotide products (Fig. 3c). The E151R variant had a limited effect on SAM-AMP synthase activity, but the D70N variant was significantly compromised, and the double mutant had no detectable activity (Fig. 3c, d). Moreover, the double mutant displayed an enhanced pppApA synthase activity when compared to the wild-type enzyme, suggesting a partial reversion of the acceptor binding site to favour ATP over SAM (Extended Data Fig. 7). A deeper understanding of the reaction mechanism and substrate specificity of the SAM-AMP synthases will require structural data in the presence of ligands, and could also involve discrimination by the Cas5 subunit, which is in the vicinity of the ATP2 ligand.

## CRISPR ancillary effectors bind and degrade SAM-AMP

To test the hypothesis that SAM-AMP is the activator of the CorA effector, we expressed *B. fragilis* CorA in *E. coli* and purified the protein to near-homogeneity in the presence of detergent (Extended Data, Fig. 8a). CorA was incubated with radiolabelled SAM-AMP, SAH-AMP or cA_3_ and then analysed by native gel electrophoresis. A clear shifted species, close to the wells of the gel, was observed to accumulate as the CorA protein was incubated at increasing concentrations with SAM-AMP or SAH-AMP (Fig. 4a). In contrast, cA_3_ was not shifted. These data support a model where CorA binds the SAM-AMP second messenger to provide immunity. Although the mechanism of the CorA effector has not yet been determined, it most likely functions as a SAM-AMP activated membrane channel, analogous to the Csx28 protein associated with Cas13 ^37^ and to a number of other predicted membrane proteins associated with type III CRISPR systems ^16^

As described previously, most type III systems with a CorA effector also encode a PDE of the NrN or DEDD family. We therefore incubated the *B. fragilis* NrN protein with SAM-AMP and observed that it specifically degrades SAM-AMP (Fig. 4b), but not the linear dinucleotide pApA, or cyclic oligoadenylate molecules cA_2-6_ (Extended Data, Fig. 8b). This suggests that specialised NrN and DEDD family PDEs may represent a type of “off-switch” to reset the system, analogous to the ring nucleases that degrade cyclic oligoadenylates in canonical type III CRISPR systems ^10^. This conclusion is supported by the observation that the CorA protein is toxic in the absence of the NrN effector (Fig. 1c). In the *Clostridia*, the NrN protein is replaced by a predicted SAM lyase (Fig. 1b), suggesting an alternative means to degrade the SAM-AMP signalling molecule. We tested this hypothesis by cloning and expressing the SAM lyase from *C. botulinum* and measuring its ability to degrade SAM-AMP, observing efficient degradation of the molecule to 5’-methylthioadenosine (MTA) (Fig. 4b, c). The other product of a lyase reaction, L-homoserine lactone (HL), is not UV visible. The *C. botulinum* lyase degrades SAM-AMP more efficiently than SAM (Extended Data, Fig. 8c), consistent with a specialised role in defence.

## Discussion

The polymerase active site of Cas10, the catalytic subunit of type III CRISPR systems, which consists of two DNA polymerase family B Palm domains, is known to synthesise a range of cyclic oligoadenylate second messengers for antiviral defence. Here, we have shown that some type III CRISPR systems signal via synthesis of a previously unknown molecule, SAM-AMP, created by the adenylation of *S*-adenosyl methionine (Fig. 5). SAM-AMP has not been described previously, either as a natural or synthetic product. As such, it represents a new type of signalling molecule, further expanding the range of nucleotide-based second messengers. In bacteria, recently discovered anti-phage signalling molecules include the cUMP and cCMP of the PYCSAR system ^38^, a wide range of cyclic di- and tri-nucleotides of the CBASS system ^39^ and the cyclic oligoadenylates typically made by type III CRISPR systems ^6, 7^. Given the structural similarity between ATP and *S*-adenosyl methionine, it is perhaps not surprising in retrospect that SAM can substitute for ATP as an acceptor for a new 5’-3’ phosphodiester bond in the active site of nucleotide cyclases, following limited sequence divergence. Clearly this reaction reaches a natural end point as there is no possibility of cyclization or further polymerization. Judging by the distribution of CorA effectors, SAM-AMP signalling has a patchy but wide distribution in members of the *bacteroidetes*, *firmicutes*, *δ-* and *ε-proteobacteria* and *euryarchaea*. This is consistent with the high levels of defence system gain (by lateral gene transfer) and loss observed generally and may be a reflection of the pressures exerted by viruses, driving diversity. Our bioinformatic analysis suggests that the ability to synthesize SAM-AMP may have arisen independently in three distinct lineages of Cas10 (Fig. 1a), perhaps in response to the prevalence of viral ring nucleases that degrade cOA. Furthermore, it is reasonable to predict that CorA is not the only ancillary effector family that is modulated by SAM-AMP, opening up new avenues for discovery.

CRISPR-associated CorA proteins are predicted to have a N-terminal soluble domain fused to a C-terminal trans-membrane helical domain related to the CorA family of divalent cation transporters ^16^. We postulate that binding of SAM-AMP to the cytoplasmic domain results in an opening of the trans-membrane pore to effect immunity, but alternative mechanisms of membrane disruption have been observed for bacterial immune effectors ^40^, so this is a priority for future studies. The CorA effectors seem to be obligately associated with degradative enzymes such as NrN in *B. fragilis*, sometimes even being fused ^16^. Our data shows that both the CorA and NrN proteins are required for defence in plasmid challenge experiments, and that without NrN the CorA protein can be toxic. The observation that SAM-AMP is easily purified from extracts of *E. coli* expressing the activated *B. fragilis* Cmr system suggests that SAM-AMP is not a substrate for the generalist ribonucleases present in bacteria, necessitating the addition of a specialised PDE such as NrN. In these respects, NrN is reminiscent of the ring nucleases (Crn1-3, Csx3) that are frequently found associated with cOA generating CRISPR systems ^10^. This suggests that it is beneficial to the cell to destroy the SAM-AMP signalling molecule, perhaps to avoid unnecessary cell death when phage infection has been cleared. In this regard, it is telling that the NrN phosphodiesterase is sometimes replaced by a SAM-AMP lyase – an enzyme that degrades SAM-AMP using an entirely different mechanism ^24^, yielding different products. SAM lyases are typically phage related genes and are thought to function by neutralising DNA methylases in host R-M systems ^24, 25^. In the context of CorA-family CRISPR systems, SAM-AMP lyases encoded by MGEs may also function as anti-CRISPRs just as viral ring nucleases do ^41^. The strict requirement for both NrN and CorA for effective immunity in the plasmid challenge assays is somewhat surprising, but should be caveated given the use of an overexpressed system in a heterologous host bacterium.

Given the wide range of AdoMet and ATP analogues available, the discovery of an enzymatic route to synthesis of SAM-AMP opens the way to the generation of a new family of bioactive molecules. For example, there is considerable interest in the development of specific inhibitors of methyltransferases, a large family of enzymes (over 300 methyltransferases encoded in the human genome) which are involved in many key cellular reactions ^42^. Depending on the specificity of the Cas10 enzyme, a range of SAM and ATP analogues could be provided as building blocks to make a diverse family of SAM-AMP analogues with altered properties. For example, replacement of the methyl group on the sulfur atom of AdoMet with a proton (i.e. S-adenosyl homocysteine) or an amino group (sinefungin) still supports catalysis by *B. fragilis* Cas10. Many other modification sites are available on the parental molecules.

In conclusion, we report the discovery of the previously undescribed SAM-AMP molecule, synthesised from two of the most abundant molecules in the cell as a second messenger of viral infection. This broadens the repertoire of type III CRISPR systems and may have implications for immune signalling more generally, as family B polymerases are a widespread and diverse superfamily found in all branches of the tree of life. The recent dramatic expansion of our knowledge of signalling molecules reflects the fact that Nature tends to use and repurpose such molecules in diverse cellular processes.

## Acknowledgements

This work was supported by grants from the Biotechnology and Biological Sciences Research Council (Grant BB/T004789/1 to MFW) and European Research Council (ref. 101018608 to MFW). HC acknowledges the support of the China Scholarship Council (code 202008420207). VH is funded by the Finnish Cultural Foundation. Thanks to David O’Hagan for helpful discussions.

## Author contributions

H.C. planned, carried out and analysed the experiments and drafted the manuscript; V.H. carried out the bioinformatic analyses; S. G. planned the protein expression and plasmid challenge experiments with H.C; S.Gra. cloned genes and expressed proteins; S.S. carried out the mass spectrometry and analysed the data; M.F.W. conceptualised and oversaw the project, obtained funding and analysed data along with the other authors. All authors contributed to the drafting and revision of the manuscript.

## Additional information

Supplementary information is available for this paper. Correspondence and requests for materials should be addressed to Malcolm White.

## Methods

### Cloning

Table S1 shows the synthetic gene, DNA and RNA oligonucleotide sequences used in this study.

The synthetic genes encoding *B.fragilis* Cas6, CorA, NrN and *C. botulinum* SAM lyase and their variants were codon optimised for expression in *E. coli* C43 (DE3) via the vector pEhisV5Tev, which encodes eight histidines, a V5 epitope tag and the cleavage site of Tobacco Etch Virus (TEV) protease ^43^.

The pACE-based Cmr expression plasmid pBfrCmr1-6 was designed to contain six codon-optimised genes *cmr1*-*6* (Twist Biosciences), with a polyhistidine tag on the N-terminus of Cmr3. We then divided pBfrCmr1-6 into five overlapping segments with similar length (designated as BfrCmr a, b, c, d and e). These segments were assembled into pBfrCmr1-6 through NEBuilder® HiFi DNA Assembly Master Kit. The obtained plasmid pBfrCmr1-6 was verified by restriction digest and sequencing.

For the construction of the CRISPR RNA over-expression vector, the codon-optimised *cas6* gene was synthesised as a g-Block (IDT) and inserted into the *Nde*I and *Xho*I restriction sites in MCS-2 of the vector pCDFDuet. The synthetic gene of CRISPR pre-array with two CRISPR repeats and two divergent *Bpi*I sites between two repeats for a spacer sequence insertion was cloned into 5’-*Nco*I and 3’-*Sal*I sites in MCS-1 of pCDFDuet containing *cas6*. Spacers targeting the tetracycline resistance gene *tetR* or *lacZ* were ligated into the *Bpi*I sites of CRISPR pre-array to obtain the plasmid, designated as pCRISPR_Tet or pCRISPR_pUC. A CRISPR array, consisting of one spacer targeting the gene encoding Late Promoter Activating protein (Lpa) of phage P1, flanked by two *B. fragilis* CRISPR repeat sequences, was assembled by annealing primers Bfr-rep-5p-T, Bfr-rep-5p-C, Bfr-rep-3p-T, Bfr-rep-3p-C, Bfr-sp-phageIPA-T and Bfr-sp-phageIPA-C (Table S1). The array was then ligated into MCS-1 of pCDFDuet containing *cas6* in MCS-2 to give pCRISPR_Lpa.

To construct pRATDuet-derived plasmids. For single effector expression, synthetic genes encoding CorA, NrN or their variants were inserted into sites of *Nco*I and *EcoR*I in MCS-1 under control of the pBAD promoter. The *nrn* gene was cloned into the *Nde*I and *Xho*I site in MCS-2 of pRATDuet containing *corA* in MCS-1 for effector co-expression.

### Protein Production and Purification

*B. fragilis* CorA, NrN, Cas6 and *C. botulinum* SAM lyase were expressed and purified using standard procedures described previously, with removal of the N-terminal his-tags by TEV protease ^43^. Subsequently, a series of variants were constructed and purified following the same procedure as the wild type. The Cmr complex with crRNA expressed from cells co-transformed with pACE-BfrCmr and pCRISPR_Lpa was purified using the same procedures.

### Plasmid challenge assay

pBfrCmr1–6 and pCRISPR_Tet (or pCRISPR_pUC) were co-transformed into *E. coli* Bl21star. Single colonies were picked for competent cell preparation into L-Broth (100 µg/ml ampicillin and 50 µg/ml spectinomycin) and cultivated at 37 °C overnight. Overnight culture was diluted 50-fold into 20 ml selective LB medium and grown at 37 °C with shaking at 220 rpm until the OD_600_ reached 0.4 - 0.5. Cell pellets were collected and then resuspended in an equal volume of pre-chilled competent cells solution (60 mM CaCl_2_, 25 mM MES, pH 5.8, 5 mM MgCl_2_, 5 mM MnCl_2_). Cells were incubated on ice for 1 h, centrifuged and the collected pellet was resuspended in 0.1 volumes of the same buffer containing 10 % glycerol. Aliquots (100 µl) were flash frozen by liquid nitrogen and then stored at −80 °C. 100 ng pRAT or pRAT derived plasmids carrying the target gene were added to the competent cells, incubated on ice for 30 min and transformed by heat shock. Following addition of 0.5 ml LB medium, the transformation mixture was incubated at 37 °C for 2.5 h. 3 µl of a 10-fold serial dilution was applied in duplicate to LB agar plates (supplemented with 100 µg/ml ampicillin and 50 µg/ml spectinomycin) for the uninduced condition. The transformants were selected on LB agar containing 100 µg/ml ampicillin, 50 µg/ml spectinomycin and 12.5 µg/ml tetracycline. Addition of 0.2 % (w/v) - lactose and 0.2 % (w/v) - L-arabinose was used for full induction. Plates were incubated at 37 °C overnight. The experiment was performed as two independent experiments with two biological replicates and at least two technical replicates.

### Synthesis of SAM-AMP and its analogues

For *in vitro* synthesis, 2 µM wild type Cmr was incubated with ATP and SAM, SAH or sinefungin (0.1 mM each for radio-labelled products or 0.5 mM each for HPLC analysis) in reaction buffer (20 mM Tris-HCl, pH 7.5, 10 mM NaCl, 1 % glycerol and 5 mM MnCl_2_). The reaction was initiated by adding 5 µM target RNA-Lpa (using non-target RNA-pUC as negative control) and carried out at 37 °C for 1 hr or the times indicated. 4 nM _α_^32^P-ATP as a tracer was added in each reaction to generate radio-labelled products, if needed for TLC analysis or EMSA.

For *in vivo* production, a single colony of *E. coli* Bl21star transformed with the plasmids pBfrCmr1-6, pCRISPR_Tet (or pCRISPR_pUC) and pRATDuet was inoculated into 10 mL of L-broth with antibiotic (50 µg/ml ampicillin, 50 µg/ml spectinomycin and 12.5 µg tetracycline) and grown overnight at 37 °C with shaking at 180 rpm. 20-fold overnight culture was diluted into 20 ml fresh L-broth with same antibiotics and then incubated at 37 °C. The culture was fully induced with 0.2 % (w/v) D-lactose and 0.2 % (w/v) L-arabinose after reaching an OD_600_ between 0.4 and 0.6. After overnight induction at 25 °C, the cell culture was mixed with 4 volumes of cold PBS and then centrifuged at 4000 rpm for 10 min at 4 °C. Cell pellet was resuspended into 2 ml cold extraction solvent [acetonitrile/methanol/water (2/2/1, v/v/v)], vortexed for 30 s and stored at −20 °C until needed. The supernatant was obtained by centrifuging at 13,200 rpm for 10 min at 4 °C, followed by evaporation until samples were completely dry, then resuspended in water and analysed by HPLC or LC-MS.

### Treatment with nuclease P1

100 _μ_M of compound was incubated with 0.02 units P1 nuclease (New England Biolabs) in P1 reaction buffer (50 mM sodium acetate pH 5.5) at 37 °C for 1 h. Each reaction solution was deproteinised with a spin filter (Pall Nanosep®, MWCO 3 kDa) followed by HPLC analysis.

### Liquid Chromatography and Mass Spectrometry

Enzymatic reactions were analysed on an UltiMate 3000 UHPLC system (ThermoFisher scientific) with absorbance monitoring at 260 nm. Samples were injected into a C18 column (Kinetex EVO 2.1 × 50 mm, particle size 2.6 µm) at 40 °C. Gradient elution was performed with solvent A (10 mM ammonium bicarbonate) and solvent B (acetonitrile plus 0.1 % TFA) at a flow rate of 0.3 ml/min as follows: 0-0.5 min, 0 % B; 0.5-3.5 min, 20 % B; 3.5-5 min, 50 % B; 5-7 min, 100 % B.

LC-MS and LC-MSMS analysis were carried out on a Eksigent 400 LC coupled to Sciex 6600 QTof mass spectrometer, in trap elute configuration at micro flow rates. Sample were loaded onto a YMC Triart C18 trap cartridge 0.5 x 5.0mm in 99.95 % water, 0.05 % TFA at 10 µl/min. After 3 min washing the salts to waste, the trap was switched in-line with the analytical column: a YMC Triart 150 x 0.075 mm. Gradient elution was performed with solvent A (99.9 % water, 0.1% FA) and solvent B (20 % water 80 % acetonitrile 0.1 % FA) at a flow rate of 5 µl/min as follows: 0 min 3% B; 0-6 min 95 % B; 6-8 min 95 % B; 8-9 min 3 % B; 9-13 min re-equilibrate 3 % B. The flow from the column sprayed directly into the ESI turbospray orifice of the MS. Data were collected in positive ionization mode from 120-1000 m/z. Ions of interest were selected for CID fragmentation at collision voltages of 25-45 V, and the fragmentation spectra collected from 50-1000 m/z. The mass spectrometer was externally calibrated prior to analysis with Sciex tuning solution 4457953.

### Thin layer chromatography

Radio-labelled SAM-AMP and its analogues were separated by TLC. 1 µl radio-labelled products were analysed on 20 × 20 cm Silica gel TLC aluminum plates (Sigma-Aldrich) with 0.5 cm of TLC buffer (0.2 M ammonium bicarbonate, 70 % ethanol, and 30 % water pH 9.3) at 35 °C. The TLC plate was left in the TLC chamber until the solvent front was approximately 5 cm from the top of the TLC plate and finally phosphorimaged using a Typhoon FLA 7000 imager (GE Healthcare).

### Electrophoretic Mobility Shift Assay (EMSA)

40 nM ^32^P-radiolabelled SAM-AMP, SAH-AMP or cA_3_ was incubated with varying amounts of purified BfaCorA (0.1, 0.2, 0.4, 0.8, 1.6 and 3.3 µM) in binding buffer (12.5 mM Tris-HCl, pH 8.0, 10 % (v/v) glycerol, 0.5 mM EDTA) at 25 °C for 15 min. Reactions were mixed with ficoll loading buffer and then analysed on the native polyacrylamide gel (8 % (w/v) 19:1 acrylamide:bis-acrylamide). Electrophoresis was carried out at 200 V for 2 hr at room temperature using 1X TBE buffer as the running buffer, followed by phosphor imaging (Typhoon FLA 7000 imager (GE Healthcare), PMT = 700-900).

### Nuclease assay

Nuclease activity of Cas6 was assayed by incubating 1.2 µM BfaCas6 with 300 nM 5’ end FAM-labelled *B. fragilis* CRISPR repeat RNA (Table S1, purchased from Integrated DNA Technologies (IDT)) in reaction buffer (20 mM Tris-HCl pH 7.5, 50 mM NaCl and 1 mM EDTA), at 37 °C for 5 min. The reaction was stopped by heating at 95 °C for 5 min and then analysed by 20 % acrylamide, 7 M urea, 1 X TBE denaturing gel, which was run at 30 W, 45 °C for 2 h. An alkaline hydrolysis ladder was generated by incubating RNA in 5 mM NaHCO_3_, pH 9.5 at 95 °C for 5 min. The gel was finally imaged using a Typhoon FLA 7000 imager (GE Healthcare) at a wavelength of 532 nm (PMT = 600-700).

An internally radio-labelled transcript RNA containing two CRISPR repeats and one guide sequence (Table S1) was incubated with 2 µM Cas6 in the same condition mentioned above and the reaction products were checked on the 20 % acrylamide denaturing gel at different time points. The transcript was generated by following the instruction of MEGAscript®Kit (Invitrogen). The template used in transcription was obtained by PCR of plasmid pCRISPR_Lpa using primer Duet-up and Duet-Down (Table S1, purchased from IDT). PCR product (120 ng) mixed with ATP, GTP, UTP, CTP solution and 133 nM _α_^32^P-ATP as a tracer was incubated at 37 °C for 4 h in the 1X reaction buffer with T7 enzyme mix. Transcript was then purified by phenol: chloroform extraction and isopropanol precipitation.

### Target RNA Cleavage Assay

5’ end labelled RNA-Lpa was generated as described previously ^15^. RNA cleavage assays using 1 µM wild type Cmr (or variant with Cmr4 D27A) and 5’ end-labelled RNA-Lpa substrates were conducted in reaction buffer (20 mM Tris-HCl, pH 7.5, 10 mM NaCl, 1 % glycerol and 5 mM MnCl_2_, 0.1 U µl^-1^ SUPERase•In™ (Thermo Scientific)) at 37 °C. The reaction was stopped at indicated time points by adding EDTA (pH 8.0) and extracted with phenol-chloroform to remove protein. After adding equal volume 100 % formamide, the samples were loaded onto 20 % denaturing polyacrylamide sequencing gel. The gel electrophoresis was carried out at 90 W for 3 – 4 h. Visualization was achieved by phosphorimaging as above. A 5’ end-labelled RNA-Lpa was subjected to alkaline hydrolysis generating a single nucleotide resolution ladder for RNA size determination.

### Bioinformatic analyses

To investigate the phylogenetic diversity of Cas10 proteins across type III CRISPR-Cas loci and their associations with CorA, all bacterial and archaeal genomes from NCBI marked as “complete” were downloaded on March 14^th^ 2023. These 51,905 genomes (50,924 bacterial and 992 archaeal genomes) were filtered for the presence of Cas10 by performing a Hidden Markov Model (HMM) search using Hmmer 3.3.2 ^44^ and previously published Cas10 HMM profiles ^18^. Hits with an E-value less than 1e-20 and minimum protein length above 500 AA were considered. The Cas10s from the resulting 2209 genomes were clustered with CD-hit (similarity cutoff 0.80 and word size 5) resulting in 613 representative Cas10 sequences. All CRISPR-Cas loci in the associated 613 genomes were annotated with CRISPRCasTyper 1.8.0 ^18^ and type III loci (or hybrid loci with type III as a component) that had an interference completeness percentage of over 30% were picked for further analysis. This threshold was chosen to exclude any solo Cas10 proteins, but also to include small CRISPR-Cas loci that may include effector proteins such as CorA. From this point forward all CRISPR-Cas loci were treated as independent of their host, i.e multiple type III loci from a single host were permitted, resulting in 745 loci. Proteins within these loci (and 2 kbp upstream and downstream of the CCTyper-reported locus boundaries) were searched for CorA using Hmmer 3.3.2 ^44^ with HMM profiles derived from CCTyper ^18^ using an E-value cutoff of 1e-20. To create the Cas10 phylogenetic tree, the Cas10 proteins from each locus were aligned using Muscle 5.1 ^45^ with the Super5 algorithm intended for large datasets. The Cas10 phylogenetic tree was constructed from the alignment using FastTree 2.1.11 ^46^ with the WAG+CA model and Gamma20-based likelihood. The tree was visualized in R 4.1.1 and RStudio 2021.9.0.351 (RStudio Team 2021. RStudio: Integrated Development Environment for R. RStudio, PBC, Boston, MA URL http://www.rstudio.com/) using ggtree ^47^ and ggplot2 ^48^. These steps were wrapped in a Snakemake 7.22.0 ^49^ pipeline and an R script that are available in Github: https://github.com/vihoikka/Cas10_prober.

## Data Availability

Mass spectrometry data are available on FigShare at the following address: <u>10.6084/m9.figshare.c.6646859</u>

---

**Extended Data Figure 1.**
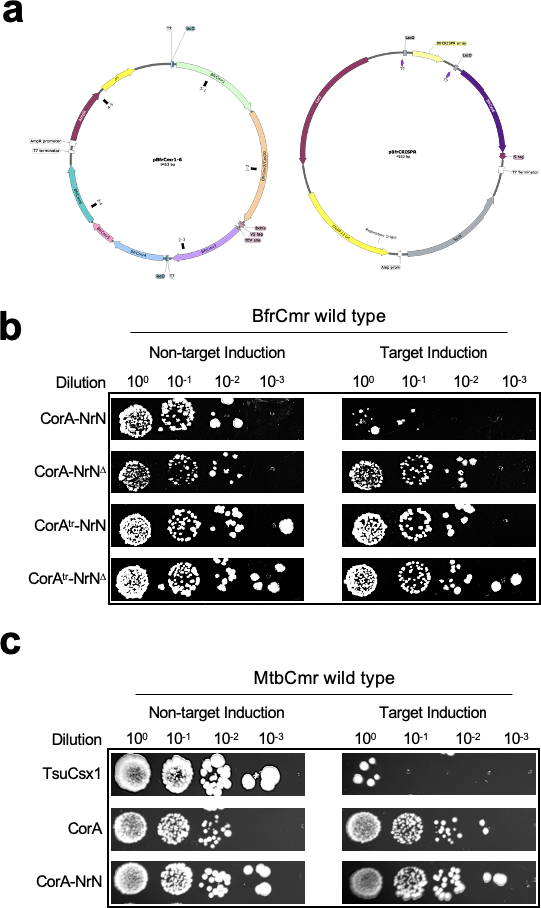
Plasmid challenge assay. **a,** Plasmid pBfrCmr1-6 was constructed by Gibson assembly to express genes cmr1–cmr6 (junctions indicated by black squares); Plasmid pCRISPR was constructed to express the CRISPR array along with Cas6. Plasmid maps were generated by SnapGene. **b,** Different variants of ancillary effectors were tested in the *B. fragilis* Cmr wild type target (tetR) and non-target (pUC19) system. CorA^tr^ is the truncated CorA (aa 1-428) and NrN^Δ^ is an inactive variant of NrN (D85A/H86A/H87A). **c,** The ancillary effectors CorA and NrN were assayed in the *Mycobacterium tuberculosis* (Mtb) Cmr wild type system. TsuCsx1 from *Thioalkalivibrio sulfidiphilus* (activated by cA_4_) acted as a positive control. Uncropped images and gels are available in Supplementary Fig. 4.

**Extended Data Figure 2.**
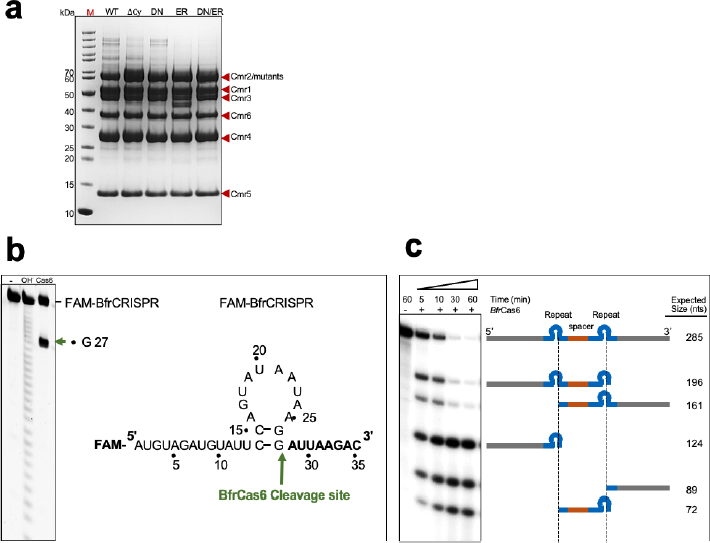
Expression of the *B. fragilis* Cmr complex and CRISPR repeat RNA processed by Cas6. **a,** SDS-PAGE analysis of the purified wild type (WT) and variants of Cmr (1-6) complex which include Cmr2 D328A:D329A (Δcy), Cmr2 D70N (DN), Cmr2 E151R (ER) and the double mutant (D70N:E151R, DNER). **b,** The cleavage site of Cas6 within the CRISPR repeat RNA was mapped by incubating 5’ end FAM-labelled repeat RNA (300 nM) with Cas6 nuclease (1.2 uM). Alkaline hydrolysis (OH^-^) ladder was used to mark the size of 5’ RNA cleavage products (green arrow). Potential secondary structure of CRISPR repeat RNA with cleavage site was indicated (green arrow). **c,** An internally radio-labelled transcript RNA containing two CRISPR repeats (blue) and one guide (targeting Phage P1) sequence (orange) was incubated with BfaCas6 (2 uM). Samples were collected at the indicated time points and analysed by denaturing gel. The expected sizes and compositions of cleavage products are indicated based on the specific cleavage site of Cas6 within each repeat. Uncropped gels are available in Supplementary Fig. 5.

**Extended Data Figure 3.**
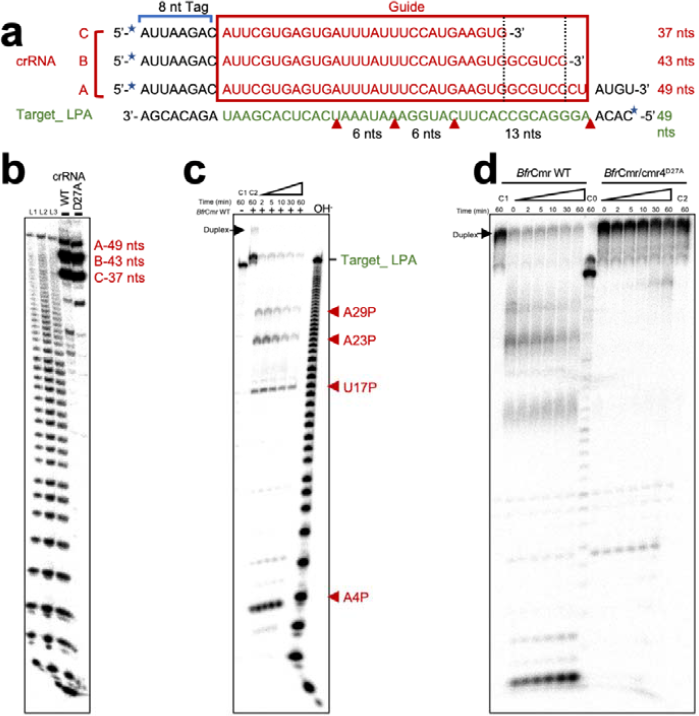
*B. fragilis* Cmr crRNA composition and target RNA degradation. **a,** The sequence of crRNA species extracted from purified Cmr and the target RNA substrate used in the activity assay. The repeat-derived sequence (8 nt tag), spacer-derived sequence (guide) and the sequence complementary to guide RNA are coloured black, red and green, respectively. Four putative cleavage sites are indicated by red arrows. Extracted crRNAs and the target RNA substrate were 5’-labelled with 32P (blue star). **b,** The size of extracted crRNAs from wild type and mutant Cmr (Cmr4D27A) was mapped by comparing with alkaline hydrolysis ladder of Target_LPA substrates (L1-3 with increased concentration of substrates). **c,** The indicated Target_LPA was incubated with (+) or without (-/C1) wild type Cmr in the presence of Mn^2+^(no Mn^2+^in buffer C2). The cleavage sites were mapped by comparing with alkaline hydrolysis (OH-) ladder and indicated by red arrow. **d,** Time course of cleavage on the 5’-radio-labelled Target_LPA by wild type or mutant Cmr (Cmr4D27A). The Buffer of C1 and C2 are in absence of Mn^2+^, while C0 is in absence of Cmr. Uncropped gels are available in Supplementary Fig. 5.

**Extended Data Figure 4.**
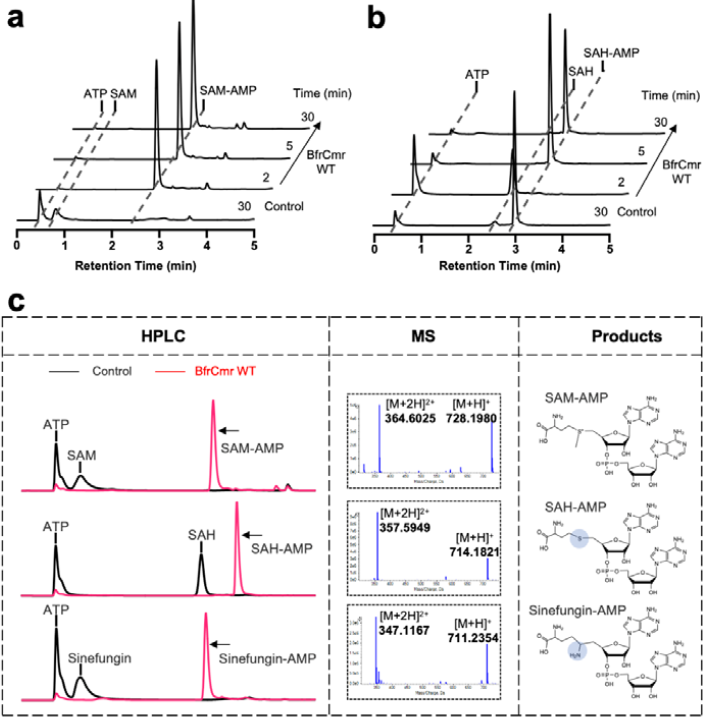
*In vitro* characterisation of *B. fragilis* Cmr-catalysed reaction. **a,** ATP (0.5 mM) and SAM (0.5 mM) were incubated with wild type of Cmr (3 µM) in presence of Mn^2+^. Samples were collected at the indicated time points and analysed by HPLC. Cmr was omitted in control samples. **b,** as for **a,** with SAH in place of SAM. **c,** HPLC and MS analysis of Cmr reaction products SAM-AMP, SAH-AMP and sinefungin-AMP. Uncropped HPLC traces are available in Supplementary Fig. 6.

**Extended Data Figure 5.**
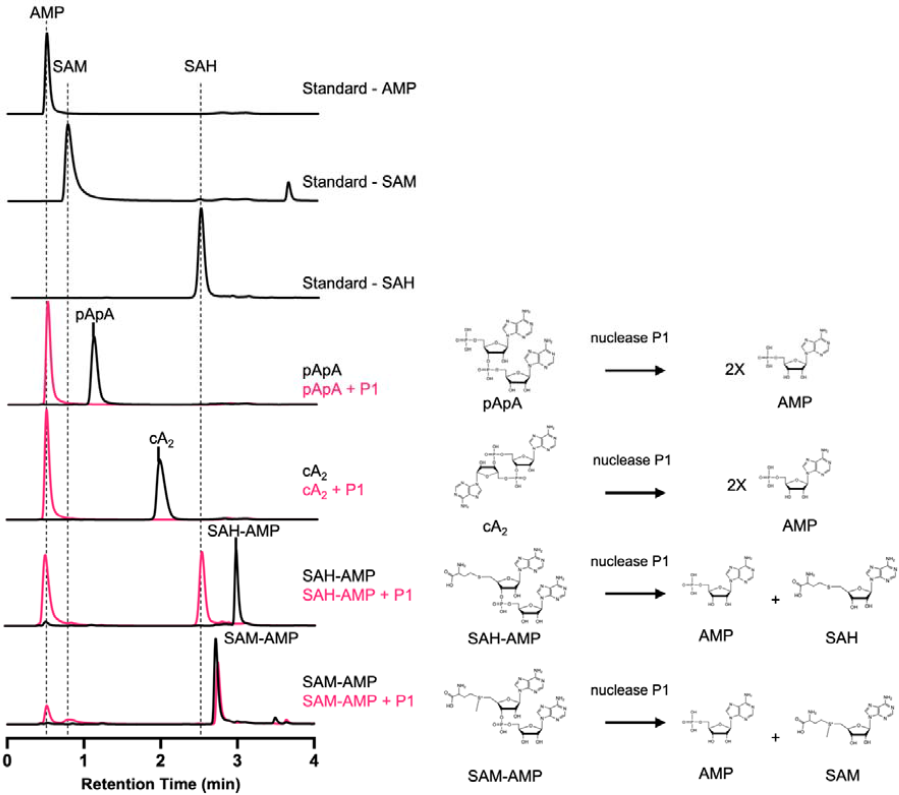
P1 nuclease degradation of reaction products. HPLC analysis of nuclease P1-mediated hydrolysis reactions towards Cmr’s products (left). cA_2_ (cyclic di-3’,5’-adenylate) and pApA (5’-phosphoadenylyl-(3’ _→_ 5’)-adenosine) were used as controls. The proposed reactions of P1 were shown on the right. Uncropped HPLC traces are available in Supplementary Fig. 6.

**Extended Data Figure 6.**
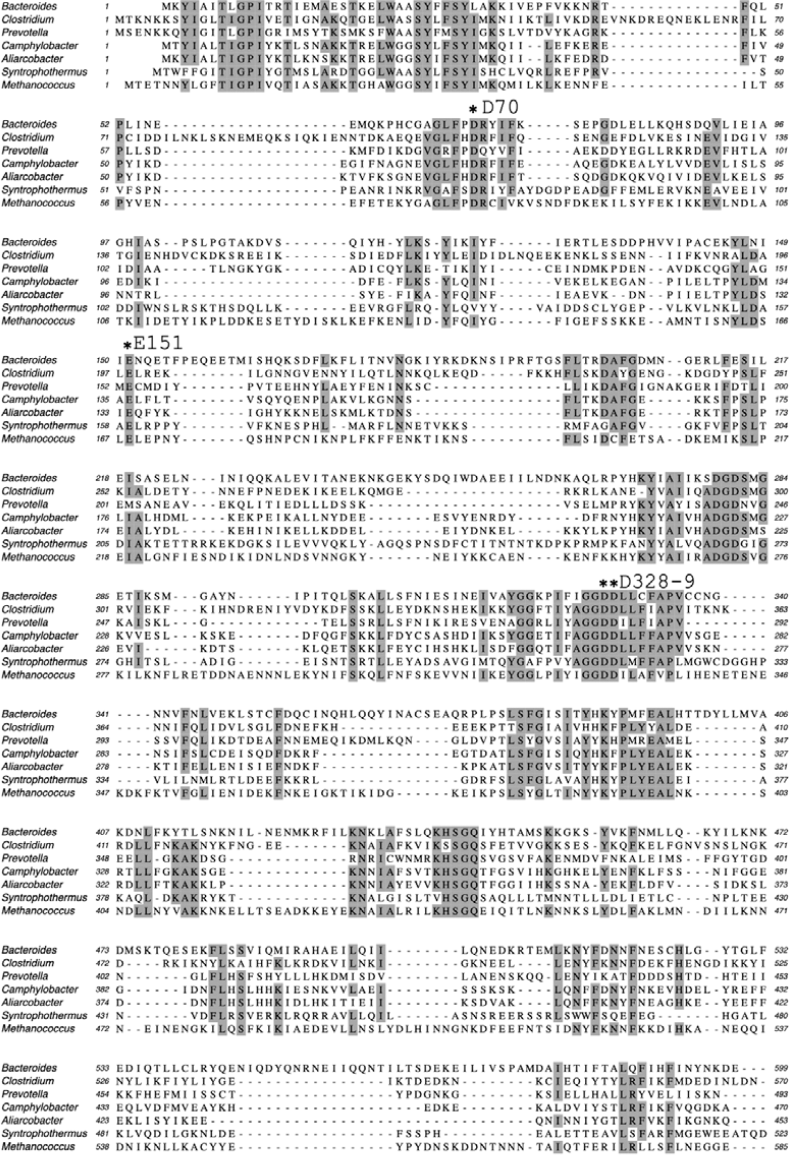
Alignment of Cas10 sequences from the CorA-1 cluster. Key binding site residues shown in figure 3 are indicated. Sequence IDs are: *Bacteroides fragilis* ANQ60746.1; *Clostridium botulinum* WP_011986674; *Prevotella - Xylanibacter muris* WP_172276208; *Camphylobacterales* bacterium HIP52383.1; *Aliarcobacter butzleri* WP_260918755; *Syntrophothermus lipocalidus* WP_013175521; *Methanococcus voltae* WP_209731901).

**Extended Data Figure 7.**
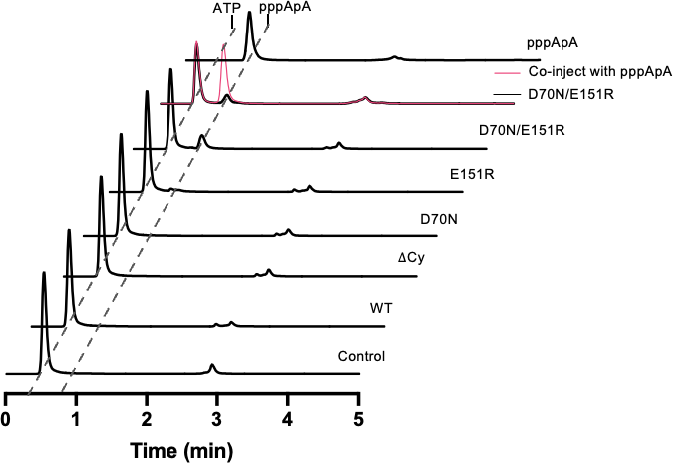
pppApA synthesis by the wild-type and variant BfrCmr complexes. *In vitro* pppApA synthase activity of wt and variant *B. fragilis* Cmr, analysed by HPLC following incubation of 2 µM Cmr with 0.5 mM ATP for 30 min. The D70N/E151R double mutant synthesises pppApA but not SAM-AMP. Uncropped HPLC traces are available in Supplementary Fig. 7.

**Extended Data Figure 8.**
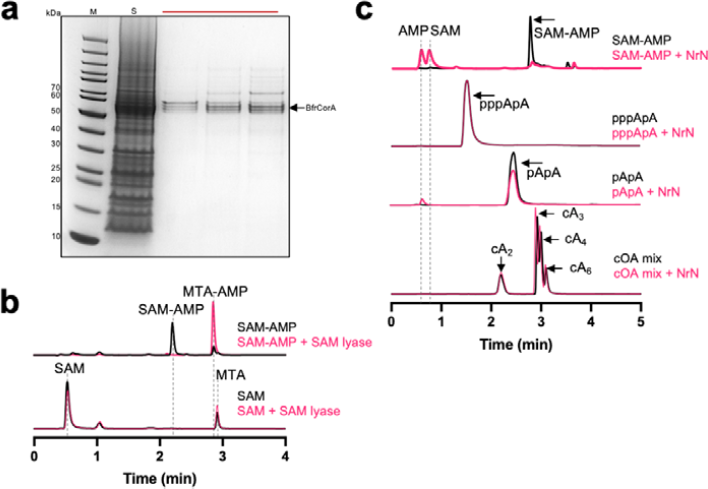
Purification of CorA and specificity of the NrN and SAM-AMP lyase effectors. **a,** SDS-PAGE analysis of the final gel filtration step for purification of *B. fragilis* CorA, which was used for SAM-AMP binding assays. The four tightly spaced protein bands in the gel all correspond to CorA, perhaps indicating limited proteolysis of the termini. M – M.Wt. markers, S – sample applied to gel filtration; the horizontal bar indicates the three fractions pooled for further analysis. **b,** HPLC analysis of reaction products for SAM-AMP and SAM following incubation with *C. botulinum* SAM-AMP lyase for 30 min. SAM-AMP is preferred as a substrate over SAM. **c,** HPLC analysis of reaction products when the NrN PDE was incubated with cA_2_, cA_3_, cA4, cA_6_, pApA and SAM-AMP for 30 min. Only SAM-AMP is a substrate for NrN. Uncropped gel and HPLC traces are available in Supplementary Fig. 8.

## Notes

### Competing Interest Statement

The authors have declared no competing interest.

